# The landscape of toxic intermediates in the metabolic networks of pathogenic fungi reveals targets for antifungal drugs

**DOI:** 10.1101/2021.09.05.459012

**Authors:** Jan Ewald, Paul Mathias Jansen, Sascha Brunke, Davina Hiller, Christian H. Luther, Humbert González-Díaz, Marcus T. Dittrich, André Fleißner, Bernhard Hube, Stefan Schuster, Christoph Kaleta

## Abstract

The burden of fungal infections for humans, animals and plants is widely underestimated and comprises deadly infections as well as great economic costs. Despite that, antifungal drugs are scarce and emergence of resistance in fungal strains contributes to a high mortality. To overcome this shortage, we propose toxic intermediates and their controlling enzymes in metabolic pathways as a resource for new targets and provide a web-service, *FunTox-Networks* to explore the landscape of toxic intermediates in the metabolic networks of fungal pathogens. The toxicity of metabolites is predicted by a new random forest regression model and is available for over one hundred fungal species. Further, for major fungal pathogens, metabolic networks from the KEGG database were enriched with data of toxicity and regulatory effort for each enzyme to support identification of targets. We determined several toxic intermediates in fungal-specific pathways like amino acid synthesis, nitrogen and sulfur assimilation, and the glyoxylate bypass. For the latter, we show experimentally that growth of the pathogen *Candida albicans* is inhibited when the detoxifying enzymes Mls1 and Hbr2 are deleted and toxic glyoxylate accumulates in the cell. Thus, toxic pathway intermediates and their controlling enzymes represent an untapped resource of antifungal targets.

## 1 INTRODUCTION

Fungi represent important human pathogens which cause a wide range of diseases from harmless superficial infections of skin and nails, affecting roughly a quarter of the worldwide population [1], to life-threatening infections with millions of patients per year. Despite lower incidence rates globally of invasive fungal infections than e.g. tuberculosis, Brown *et al.* [2] estimates a comparable number of deaths (around 1.5 millions) due to the very high mortality rates of invasive fungal infections. For example, in France the total burden of serious fungal infections is estimated to affect more than 1% of the population each year [3] and the current COVID-19 pandemic is accompanied by life-threatening secondary fungal infections, predominately by SARS-CoV-2–associated pulmonary aspergillosis (CAPA) [4] or mucormycosis [5].

The increasing incidence of invasive fungal infections is linked to advances in medical care that can save lives of immunocompromised patients, which however are at high risk of fungal infections [6, 7]. While most immunocompetent hosts are able to clear or control fungal infections, immunocompromised patients can suffer from blood-borne disseminating infections with often very high mortality rates of more than 50 % [2]. However, there is only a very limited number of antifungal agents. This is mostly due to the much closer phylogenetic relationship between fungi and humans compared to bacterial pathogens, which constrains the options for agents that do not harm the human host [8]. Furthermore, resistance to the widely used azole antifungals is increasingly reported [9] and polyenes antifungals are only cautiously used due to their toxicity to mammalian cells [9]. Due to these difficulties, novel approaches for speeding up the discovery of antifungal drugs and their targets are urgently needed [10, 11].

The metabolic flexibility of pathogenic fungi is a cornerstone of their virulence [12] and, additionally, enzymes of fungal-specific pathways in central metabolism such as ergosterol biosynthesis are key targets for antifungal drugs [10, 11, 13]. Hence, the metabolism of pathogenic fungi has come under scrutiny to establish novel antifungal targets and develop highly efficient new antimycotics [14, 15]. Since the metabolism of fungi is highly dynamic during host interactions [12, 16], modeling of metabolic regulation by dynamic optimization is of high relevance to unveil optimal regulatory strategies in fungal pathways as well as to elucidate key enzymes regulating pathway flux [17]. In recent years, we and others have uncovered optimality principles behind fast pathway activation strategies and efficient pathway regulation across a wide range of bacteria [18–24], which are in principle transferable to eukaryotes such as fungi, too. To combat pathogens, our previous observation that enzymes upstream of toxic intermediates are tightly regulated is of special interest, since we hypothesized that an upregulation (downregulation) of key enzymes regulating flux before (after) a toxic intermediate can lead to self-poisoning [23]. Further, as shown across prokaryotes [23], the inference of metabolic hubs and key regulated enzymes by estimators like the promoter length can be used in a reverse approach to identify highly regulated enzymes and to support target identification for potential antimicrobials. This reasoning provides a new avenue for the identification of drug targets using endogenously produced cytotoxins. While metabolic networks have recently been investigated extensively to find key enzymes for virulence using the concept of elementary modes [25], there has been, to our knowledge, no systematic screen for toxic intermediates in metabolic networks of fungal pathogens so far. However, the importance of toxic intermediates has been recognized in the fields of pathway evolution [26] and metabolic engineering [27]. Similarly, an anti-cancer therapy using endogenous toxic metabolites to stop cell growth was recently proposed [28].

For a large-scale prediction of toxicity across multiple organisms, quantitative structure - activity relationship (QSAR) models have proven to be valuable in *in silico* screening of inhibitors [29, 30], in improving yield by identification of toxic intermediates [31, 32] and in risk assessment [33, 34]. These machine learning approaches can help to fill the gap between empirical measured toxicity data in databases like ChEMBL [35] and pathway intermediates which have unknown toxicity. For fungi, Prado-Prado *et al.* [36] developed a multi-target spectral moment QSAR to classify drugs with activity against several species. However, to take advantage of novel machine learning approaches as well as much larger toxicity assays stored in the ChEMBl-database, we here used a random forest regression model to predict the toxicity of intermediates in fungal metabolism based on chemical features. Using a multi-output model [37, 38], we were able to include data from all fungi with toxicity measurements as well as different types of measurement of toxicity to train a single regression model for different fungal species and additional models for host organisms. Moreover, we integrated our predictions with data on the degree of regulation of individual enzymes in metabolic networks derived from KEGG [39]. To this end intra-species protein-protein interactions networks were inferred using the newly developed FungiWeb database (https://fungiweb.bioapps.biozentrum.uni-wuerzburg.de) and used here to overcome the lack of regulatory data like large-scale transcription factor networks in fungal species. The enriched networks together with a cytotoxicity prediction can be displayed in a web-service *FunTox*-*Networks* (http://funtox.bioinf.uni-jena.de).

As proof-of-principle for the use of our tool to identify targets for the enrichment of antifungal intermediates, we used *FunTox*-*Networks* to identify several toxic intermediates in fungal-specific pathways. As promising targets, we identified toxic intermediates and their controlling enzymes in pathways of nitrogen and sulfur assimilation, in amino acid synthesis, and in the catabolism of fatty acids as alternative carbon sources. Especially for the latter, where glyoxylate is a toxic intermediate, we show that detoxifying enzymes like malate synthase (Mls1) and an aminotransferase (Hbr2) are potential targets for novel antifungal drugs.

## 2 RESULTS

### 2.1 Prediction of metabolite toxicity with a random forest regression model

Based on the toxicity assay data in the ChEMBL database, we trained separate QSAR regression models for the prediction of metabolite toxicity in fungal species and their human as well as murine host (see section 4.1 for details). After carefully filtering wrong annotations and removal of outliers the data set for fungal species included 122,474 data points for 112 fungal species, 653,035 data points for human cells and 10,703 data points for mice (see section 4.1 for details). Due to the best performance among other regression models on a subset of the data, the random forest regression approach was chosen to build the QSAR models (see Supp. 6). The final random forest regression model for fungal species obtained a reasonable coefficient of determination of *R*^2^ = 0.64, which is slightly lower for the models of human (*R*^2^ = 0.57) and murine (*R*^2^ = 0.59) cells. The model quality is underlined by low root mean squared error (RMSE) values between 0.30 (fungi), 0.30 (human), and 0.32 (mice). For our fungal model we fulfill the recommended criteria [40] of a high *R*^2^ > 0.6 and a low RMSE, which should be lower than 10 % of the range of predicted toxicity values (between −1.73 and 4.59 in training data). Other QSAR toxicity regression models achieve higher predictive power but lack universality by considering only a single species like *Escherichia coli* [32] or by focusing on a certain group of substances with even more precise predictions [41].

While our random forest regression models show good performances on test and training data, we further used annotated compounds from the KEGG database to validate the predictions of our models. To this end, we predicted the toxicity of compounds with biological roles like antibiotics or carbohydrates in important fungal species and host organisms (see Fig. 1A). As expected, antifungal compounds were predicted to be significantly more toxic than non-antibiotic compounds in fungal species (Wilcoxon rank-sum test *P* < 1^−16^). The regression model clearly discriminates antifungals also from other antibiotics like antibacterials (Wilcoxon *P* < 11^−16^). One should note that toxicity of antibiotics is relatively high even in hosts, which is reasonable since many antibiotics cause side effects harming human cells and therefore are often not applied in high concentrations [42, 43]. Our model predicts carbohydrates as the least toxic compound group in fungi and other organism which is in line with their common biological functions and their high abundance in standard growth media and diet. From these results we conclude that our QSAR model is able to estimate the toxicity of metabolites across a wide range of compound classes and organisms.

**FIGURE 1.**
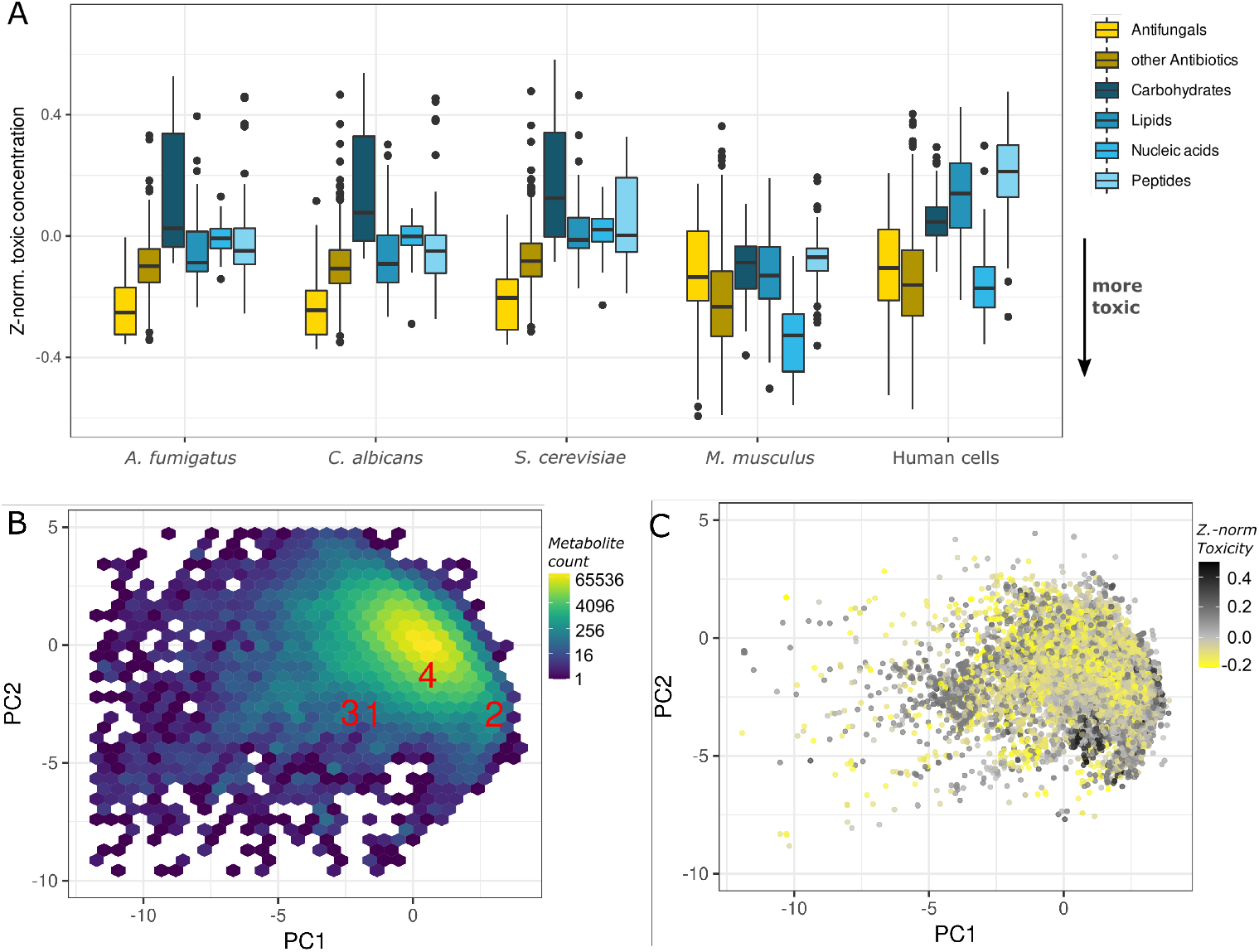
**A** Toxicity prediction of KEGG compounds and their categories in fungi and host organisms. Distribution of toxic concentration are depicted in yellow for antibiotics and in blue for common metabolites. **B** Applicability Domain of the QSAR model based on a PCA of molecular descriptors (descriptor space) with the exemplary metabolites (red): ATP (1), glyoxylate (2), amphotericin B (3), fluconazole (4). The density of learning data in descriptor space is depicted from dark blue to yellow. **C** Distribution of metabolites from KEGG based in the same descriptor space as in A and toxicity in *C*. *albicans* indicated in gradient from yellow (toxic) to black (non-toxic).

A main goal behind our QSAR model is that it is accessible for all researchers and we therefore provide the web-service *FunTox*-*Networks*. In addition to the toxicity prediction of KEGG metabolites, the web-service can be queried to predict toxicity for other or novel compounds via their Simplified Molecular Input Line Entry Specification (SMILES) or International Chemical Identifier (InChI) representation [44, 45]. Furthermore, we provide, as recommended by the Organisation for Economic Co-operation and Development (OECD) [46], information about the confidence of predictions as an applicability domain of our random forest regression model. Firstly, we provide the standard deviation across predictions of all trees as additional output, which has been shown to reflect the accuracy of predictions [47]. Secondly, a principal component analysis of trained metabolites and their molecular descriptor values was performed to visualize the descriptor space of training data (see Fig. 1B). Using this presentation we see that training data covers not only the known toxic antifungals like fluconazole or amphotericin B well, but is also nearly identical to the descriptor space obtained from all metabolites of pathways in KEGG (see Fig. 1C).

Importantly, although the toxicity prediction is mainly based on molecular descriptors, we did not observe distinct regions or clusters in the principal component analysis of molecular descriptors for toxic and non-toxic metabolites. This observation indicates that toxic metabolites are uniformly distributed across the entire chemical space (see Fig. 1C). Moreover, this demonstrates that our random forest regression model is able to recognize small structural differences that are associated with toxicity. An example is the well known toxic byproduct methylglyoxal (C3H4O2) formed during glycolysis [48]. Our model correctly predicts a high toxicity of methylglyoxal e.g. in the common fungal pathogen *Candida albicans* (*Z_tox_* = −0.177) and no toxicity of the structurally highly similar metabolite pyruvate (C_3_H_4_O_3_, *Z_tox_* = 0.007).

### 2.2 Integration of intermediate toxicity and enzyme regulation in pathway maps

Since we demonstrated a close relationship between toxicity of pathway intermediates and optimal points of pathway regulation in our previous work [23], we integrated metabolite toxicity and regulatory effort on KEGG pathway maps to identify suitable targets of deregulation to accumulate self-poisoning intermediates in pathogenic fungi (see section 4.2 and Fig. 5B for details). Thus, we provide enriched KEGG pathway maps of seven major fungal pathogens or model organisms and their hosts (human cells and mice, see Tab. 2). To infer key regulated enzymes with toxic intermediates, we used the number of transcription factors controlling an enzyme and, for pathogenic fungi, a score representing the connectivity of an enzyme within its intraspecies protein-protein interaction (PPI) network. In those networks each interaction is further characterized by its confidence, e.g. experimental validation (see section 4.2). Interestingly, yeast data show that this measure correlates with the number of interactions as well as with the number of post-translational modification (PTM) sites of a gene (*ρ* = 0.52 and *ρ* = 0.24, respectively; Spearman correlation with each *P* < 1^−16^, see also Supp. 8). Since PPI networks have not been reconstructed for all fungi, we used the promoter length as easily calculable (intergenic distance) estimator to infer transcriptional regulation in the case of *Arthroderma benhamiae* and *Aspergillus flavus* (see section 4.2).

In addition to enriched KEGG pathway maps, *FunTox*-*Networks* provides tables containing all pathways and compounds that can be searched and sorted by key features. These overview tables also show that very diverse compound groups within the KEGG pathway maps are predicted to be highly toxic. Examples in *C*. *albicans* include heme-like compounds (siroheme, precorrin), activated fatty acids (acyl-CoA) and highly reactive acids like acetic acid, which all are known to be cytotoxic [49–51]. These data further emphasize the predictive quality of our model. Examples of KEGG pathways which contain many toxic intermediates are fatty acid metabolism (map01040, map00062, map00071), propanoate metabolism (map00640), glyoxylate metabolism (map00630), and porphyrin metabolism (map00860).

Because we can predict toxicity in fungal species as well as for their potential hosts, our web-service additionally provides an interactive plot to compare the toxicity of KEGG metabolites in two species. This enables the identification of metabolites which are more toxic to the pathogen than to host cells or *vice versa* (see Fig. 2). While the former metabolites are interesting antifungals that are tolerable by the host, the latter can comprise potential small-molecule virulence factors of pathogens. Due to the advent of large-scale transcriptomics able to generate measurements during host-pathogen interaction (e.g. dual-species RNA-Seq of host and pathogen in parallel), this plot of species-specific toxicity can be enriched with previously published expression data for genes coding for the adjacent enzymes [16, 52, 53].

**FIGURE 2.**
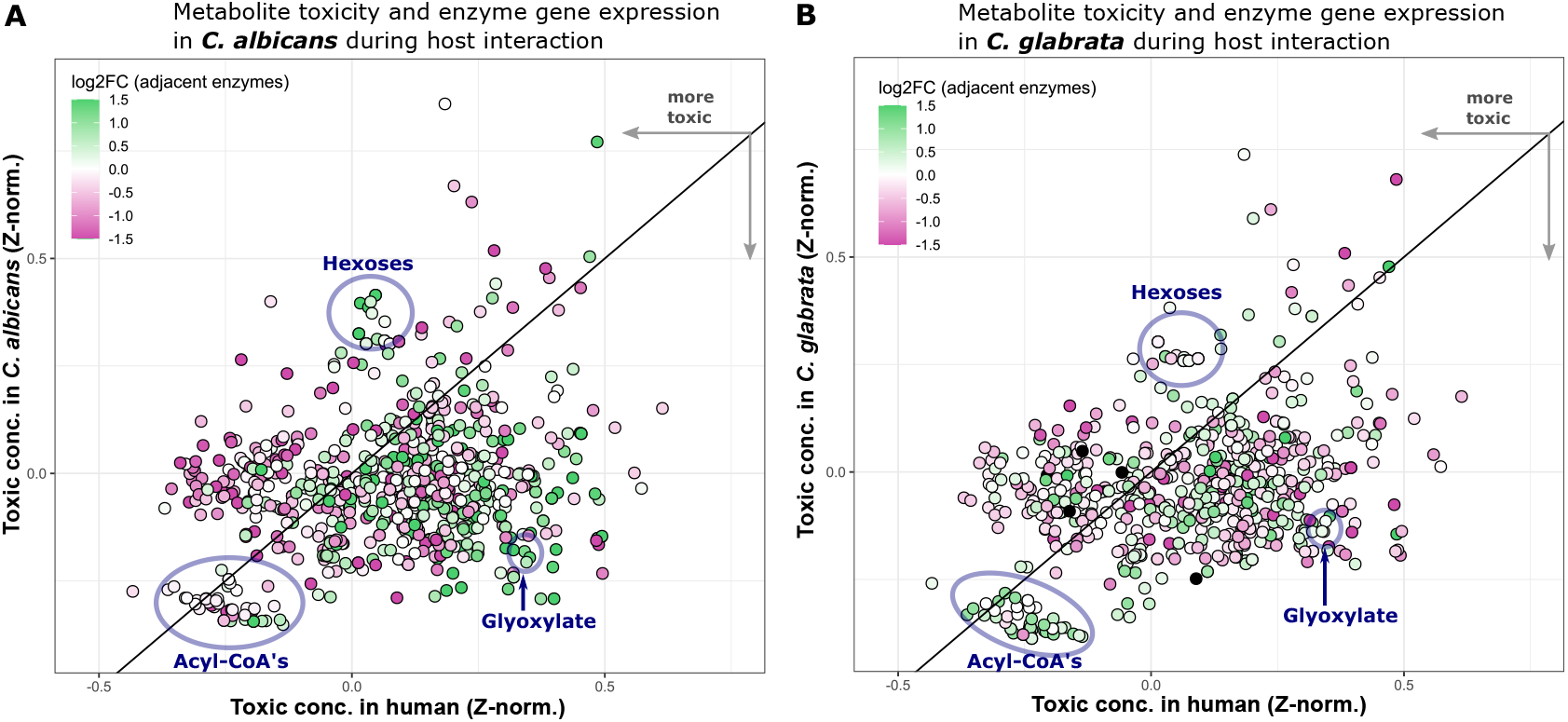
Integration of enzyme gene expression and metabolite toxicity in pathogenic fungi upon interaction with human blood cells [16]. Each dot represents the predicted toxicity of KEGG metabolites in the human host (x-axis) and in the fungal pathogen *C*. *albicans* (subpanel **A**) or *C*. *glabrata* (subpanel **B**). The color shows the up- or downregulation (green to pink) of adjacent enzyme coding genes (producing or consuming the metabolite) after 1h incubation with human blood for each *Candida* species as measured by Kämmer *et al.* [16].

Fig. 2 depicts how this view of data can be used to get insights into molecular host-pathogen interactions. Firstly, a distinct group of acyl-CoA compounds is predicted to be toxic in the pathogens *C*. *albicans* (Fig. 2A) and *C*. *glabrata* (Fig. 2B) as well as in the human host. However, the adjacent enzyme coding genes, which are mainly related to fatty-acid oxidation, are upregulated in *C*. *glabrata*, but not in *C*. *albicans* after 1h of co-incubation with human blood. In contrast, genes encoding the enzymes of the glyoxylate bypass are strongly upregulated in *C*. *albicans* and not in *C*. *glabrata*. Further, toxicity of glyoxylate is primarily predicted in fungal cells and not in host cells. Interestingly, despite the upregulation of genes encoding the enzymes of the glyoxylate bypass in *C*. *albicans*, which indicates glucose starvation, a group of non-toxic hexoses are connected to upregulated enzyme coding genes or transporter genes in sugar metabolism (see Fig. 2A).

To summarize, our toxicity prediction and web-service *FunTox*-*Networks* offers not only the prediction of toxicity for pathway intermediates, but also a detailed integration with the pathway topology in KEGG maps, regulation of enzymes, respectively the expression of their genes during host-pathogen interactions. This allows to identify new potential targets for antifungals by searching for toxic pathway intermediates and the identification of key regulated enzymes controlling the intermediate’s accumulation.

### 2.3 Toxic intermediates in fungal specific metabolic pathways

To prove its usefulness, we employed *FunTox*-*Networks* to search for enzymes which are suitable targets for antifungal interventions. We limited our search to metabolic pathways which are specific to fungi and enzymes which have no homolog in humans. Since fungal species are able to grow in a great variety of conditions, many fungal specific metabolic pathways with toxic intermediates are related to resource acquisition, such as carbon, nitrogen, and sulfur assimilation (see Fig. 3). We found known and new targets in the synthesis of amino acids and in fatty acid metabolism, as well as in ergosterol synthesis, which is a main target for the antifungal class of azoles.

**FIGURE 3.**
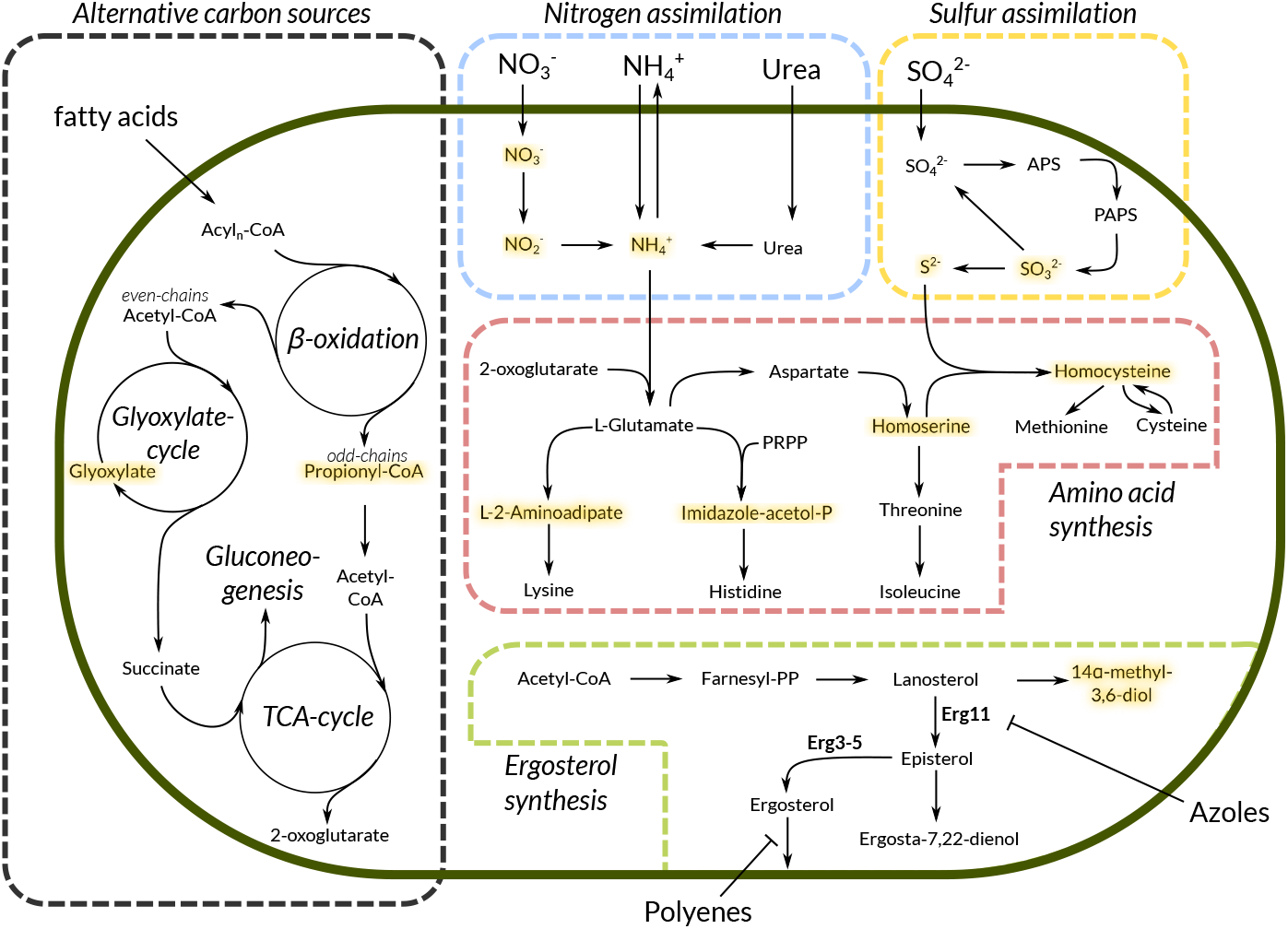
Schematic overview of target pathways with selected toxic intermediates (yellow) in a fungal cell (dark green).

Ergosterol synthesis is especially suitable for antifungal intervention in our sense, as azoles are not only inhibiting the synthesis by blocking the Lanosterol-14*α*-demethylase, but also lead to the accumulation of toxic 14-methyl sterol intermediates like 14*α*-methyl-3,6-diol [54].

To harbor suitable antifungal targets a pathway needs to be active during host invasion and is ideally essential for virulence. Because pathogenic fungi have different strategies to invade the host and conquer different host niches like the lung, gut or skin [55, 56], the importance of metabolic pathways for virulence varies across fungal species. For the target pathways shown in Fig. 3 we found that the aspartate as well as the methionine-branch of amino acid synthesis is linked to virulence [57] and especially homocysteine is known to accumulate to toxic levels if the methionine synthase gene (*MET6*) is deleted in *C*. *albicans* [58]. In contrast, the importance of lysine and histidine biosynthesis for fungal virulence is unclear due to contradictory results in common fungal pathogens [57]. Interestingly, the acquisition of inorganic nitrogen and sulfur sources is not required for full virulence and assimilation of these elements is possible via multiple organic and inorganic substrates in pathogenic fungi like *A. fumigatus* or *C*. *albicans* [59–61]. However, a recent study shows that sulfite detoxification is actively controlled and enhances growth in *C*. *albicans* suggesting that the involved enzymes and regulators are suitable targets for antifungal interventions [62].

In these metabolic pathways we considered the toxic intermediate glyoxylate with its consuming enzyme malate synthase to be the most promising target. Firstly, glyoxylate is known to be highly reactive, and eukaryotic cells isolate the glyoxylate cycle, like other pathways producing reactive species, in special compartments (peroxisomes) [63]. Additionally, the use of fatty acids as carbon sources and the use of the glyoxylate cycle as a variant of the tricarbolic acid cycle (TCA), which also allows gluconeogenesis, is a known virulence factor in *C*. *albicans* [64]. It has also been shown earlier that the enzymes unique to the glyoxylate cycle, isocitrate lyase (Icl1) and malate synthase (Mls1), are highly abundant during both, confrontation with macrophages and blood infections [16, 52, 65]. As a final line of evidence, the deletion of the malate synthase (gene *glcB*) in *Mycobacterium tuberculosis* is much more efficient in impeding the growth than deletion of the isocitrate lyase (genes *icl1* and *icl2*) due to glyoxylate toxicity [66]. This means that the glyoxylate bypass catalyzed by Icl1 (upstream of glyoxylate) and Mls1 (downstream, see Fig. 4A) can be deactivated by the inhibition of both enzymes, but the more effective antibacterial, potentially also antifungal, intervention is the inhibition of Mls1 since the toxic intermediate glyoxylate is accumulated.

**FIGURE 4.**
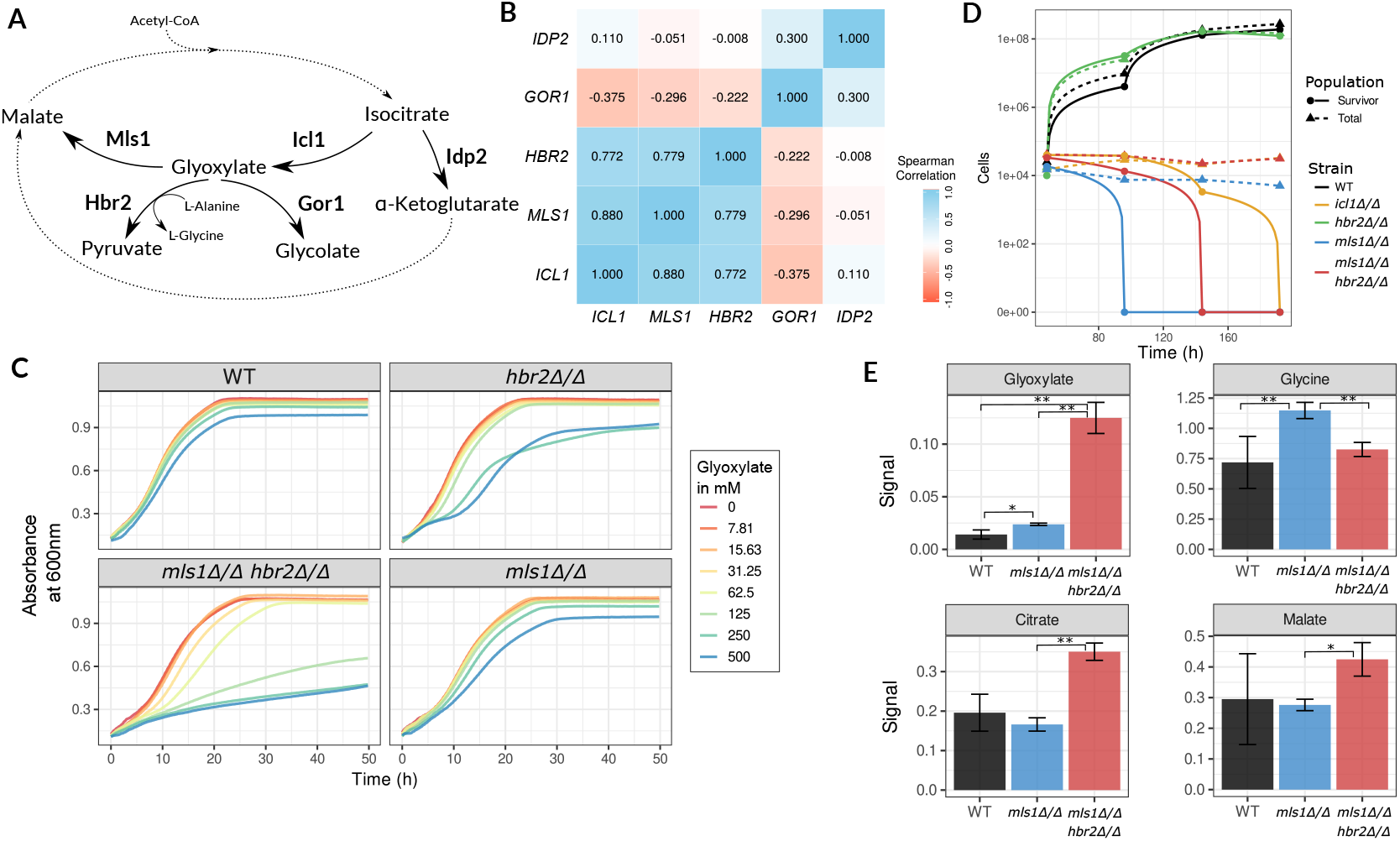
Investigation of glyoxylate detoxifying enzymes in *C*. *albicans*. **A** Overview of glyoxylate bypass as well as related metabolites and enzymes. **B** Correlation matrix of transcription for glyoxylate bypass genes (*ICL1*, *MLS1*), genes for potential detoxifying enzymes (*HBR2*, *GOR1*) and for the decarboxylation of isocitrate (*IDP2*) in response to various organic acids [67]. **C** Growth curve of *C*. *albicans* and deletion strains in glucose (2%) media with addition of varying amounts of glyoxylate. The mean of the absorbance at 600nm of three biological replicates is shown for each glyoxylate concentration. **D** Long term survival assays of *C*. *albicans* strains in SD medium and ethanol as sole carbon source. Surviving yeast cells were determined on YPD agar plates after the indicated incubation times. **E** Measurement of metabolites by gas chromatography–mass spectrometry (GC-MS) of *C*. *albicans* strains grown in YNB broth without amino acids and 2% glucose. Error bars indicate the standard errors of mean (SEM) for three replicates and significance (T-tests) is shown for selected comparisons between strains with values \**p* < 0.05, \*\**p* < 0.01, \*\*\**p* < 0.001. In all figures WT refers to the wild-type strain BWP17+CIp30.

### 2.4 The glyoxylate bypass as primary example for toxic intermediates and drug target search

To verify the potential of toxic intermediates as antifungal drug targets, we experimentally investigated the glyoxylate bypass in *C*. *albicans*. In addition to the two key enzymes Icl1 and Mls1, the genome annotation as well as metabolic databases like KEGG suggest two additional enzymes which can detoxify glyoxylate (see Fig. 4A). We therefore analyzed the transcriptional response of *C*. *albicans* to various organic acids [67]. In addition to *ICL1* or *MLS1*, we created a deletion strain lacking the putative alanine glyoxylate aminotransferase gene (*HBR2*), because it shows a strong coregulation with the glyoxylate bypass enzymes (see Fig. 4B) and its homolog in baker’s yeast codes for an enzyme that is able to detoxify glyoxylate to pyruvate [68]. In contrast to this, the glyoxylate reductase gene (*GOR1*) is coregulated with the isocitrate dehydrogenase gene *IDP2* of the TCA cycle (see Fig. 4A and B). Hence, we excluded Gor1 as candidate for the control of glyoxylate accumulation when glyoxylate bypass enzymes are active.

To study the importance of glyoxylate and their detoxifying enzymes, we performed growth experiments in glucose-rich medium at a low pH to ensure membrane-permeability of externally supplemented glyoxylate (*cf.* section 4.3). While externally added glyoxylate leads only to minor growth inhibition of the wild-type *C*. *albicans* strain, single deletion strains of *MLS1* or *HBR2* show intermediate, and a double deletion mutant (*mls1*Δ/Δ *hbr2*Δ/Δ) a significant growth inhibition (see Fig. 4C). This supports the view that glyoxylate detoxification is ensured by both, Mls1 and Hbr2.

*C*. *albicans* faces a glucose-poor environment during phagocytosis and relies on the glyoxylate bypass for survival. Having shown that *C*. *albicans* depends on Mls1 and Hbr2 for glyoxylate detoxification, we performed additional experiments where glyoxylate is produced intracellularly. To this end, we tested growth with ethanol as the only carbon source. As expected, the wild-type and *hbr*2Δ/Δ strains grew slowly on ethanol, and the glyoxylate bypass mutants were unable to grow. In addition to growth inhibition, we observed that strains lacking the glyoxylate detoxification enzymes (*mls1*Δ/Δ and *mls1*Δ/Δ *hbr2*Δ/Δ) die off earlier when ethanol is the sole carbon source than the non-glyoxylate producing strain *i cl* 1Δ/Δ, which otherwise is similarly unable to grow (see Fig. 4D).

Lastly, we confirmed glyoxylate accumulation in our mutant strains by gas chromatography–mass spectrometry (GC-MS) after growth in glucose-rich medium. We found that glyoxylate accumulated in *mls1*Δ/Δ and reached even higher concentrations in the double mutant *mls1*Δ/Δ *hbr2*Δ/Δ. This is accompanied by the enrichment of the TCA cycle intermediates citrate and malate. Interestingly, in the single mutant *mls1*Δ/Δ glycine accumulated which supports our hypothesis that glyoxylate is detoxified by Hbr2 via the conversion of alanine to glycine (see Fig. 4A).

From the above experiments we can conclude that glyoxylate accumulation cannot be achieved in *C*. *albicans* by inhibition of only the malate synthase, as it has been shown for *M. tuberculosis* [66]. Moreover, we discovered that the so-far uncharacterized aminotransferase Hbr2 is part of the more complex glyoxylate detoxification in *C*. *albicans*, which provides new avenues for drug target search. Overall, we show that toxic intermediates are an untapped resource for the development of antifungal drugs by the characterization of enzymes as well as the underlying transcriptional regulation, which control accumulation of pathway intermediates during host-pathogen interactions.

## 3 CONCLUSION AND OUTLOOK

In summary, we were able to build a new QSAR model for the prediction of metabolite toxicity for fungal species and also for their hosts during infection. Based on a specialized normalization scheme, our random forest regression models show a good performance in predicting the toxicity of different compound categories such as toxic antifungals or non-toxic carbohydrates. Since it was our goal to support antifungal drug target identification, we interleaved metabolite toxicity prediction with metabolic pathway maps for main fungal pathogens and their hosts and provide it as the web-service *FunTox*-*Networks*. To test our approach with a real-life application, we used this new resource and identified multiple potential toxic intermediates in fungal-specific metabolic pathways. We found a very promising target in the malate synthase and the aminotransferase Hbr2 of *C*. *albicans* which synergistically control the detoxification of glyoxylate. This can become important when fatty acids are used as a carbon source by *C*. *albicans* e.g. during the confrontation with phagocytic immune cells [69]. In our experiments we observed a growth defect of deletion strains related to the accumulation of toxic glyoxylate. Therefore we conclude that antifungal drugs inhibiting malate synthase and the aminotransferase Hbr2 would be more effective than inhibitors of the isocitrate lyase which are currently being tested [70]. Moreover, it reveals that a comprehensive study of the regulatory circuits controlling pathway intermediate accumulation provides an untapped resource for the discovery of novel types of antifungals.

While we focused here on fungal pathogens of humans, fungi that infect plants and animals are a significant concern in agriculture and biodiversity, leading not only to high economic costs but also affecting human health, since they can cause crop failures and famine [71]. The principle of toxic intermediates as guides to valuable antifungal drug targets can also be applied to fungal plant pathogens. Toxicity prediction via *FunTox*-*Networks* can therefore become a valuable resource for research and drug development in this field, too. Beyond the use for antifungal target selection, the prediction of intermediate toxicity can also be of advantage in industrial processes involving yeasts, which try to optimize yield and efficiency by reducing the accumulation of toxic intermediates [72]. Especially in metabolic engineering or synthetic biology, where foreign or redesigned metabolic pathways are incorporated into yeast, the knowledge of potential toxic intermediates and their control can be of great importance [73]. Interestingly, a rewiring of the glyoxylate bypass regulation was necessary to improve the titer reached in production of glycolic acid which is an important chemical compound and can be produced in yeasts by glyoxylate reduction [74, 75]. However, production is hampered by the tight repression of the isocitrate lyase when glucose is present and glyoxylate accumulation is avoided rigorously which underlines the importance of our findings.

Taken together, knowledge about toxicity of pathway intermediates as well as their regulation enables various applications in antifungal drug target search or industrial use of fungal species.

## 4 MODEL AND METHODS

### 4.1 Toxicity prediction

To train the regression model for predicting the toxicity of metabolites in fungi as well as their hosts (humans, mice), we retrieved the corresponding taxonomy-based activity data listed in the ChEMBL database [35] (Release 27, January 2021). The three data sets for fungi, human and mice were then filtered and checked for the correct taxonomy, target type (only ‘organism’ or ‘cell’ and not ‘protein inhibition’ etc.), non-valid data, only precise toxicity measurements (’=’ relation) and uniform direction of toxicity standard types (lower value indicating higher toxicity). To learn the regression model, we used a cut-off of at least 50 activity data measurements for each standard type (type of toxicity measure) and at least 100 measurements for each organism. Using these filtering steps, we obtained 122,474 activity measurements for fungi and 653,035 for humans and as well as 10,703 for mice, respectively, which were used as input for the machine learning approach (see Fig. 5). This machine learning pipeline and data is also documented and stored in a open repository (http://doi.org/10.5281/zenodo.3529162).

**FIGURE 5.**
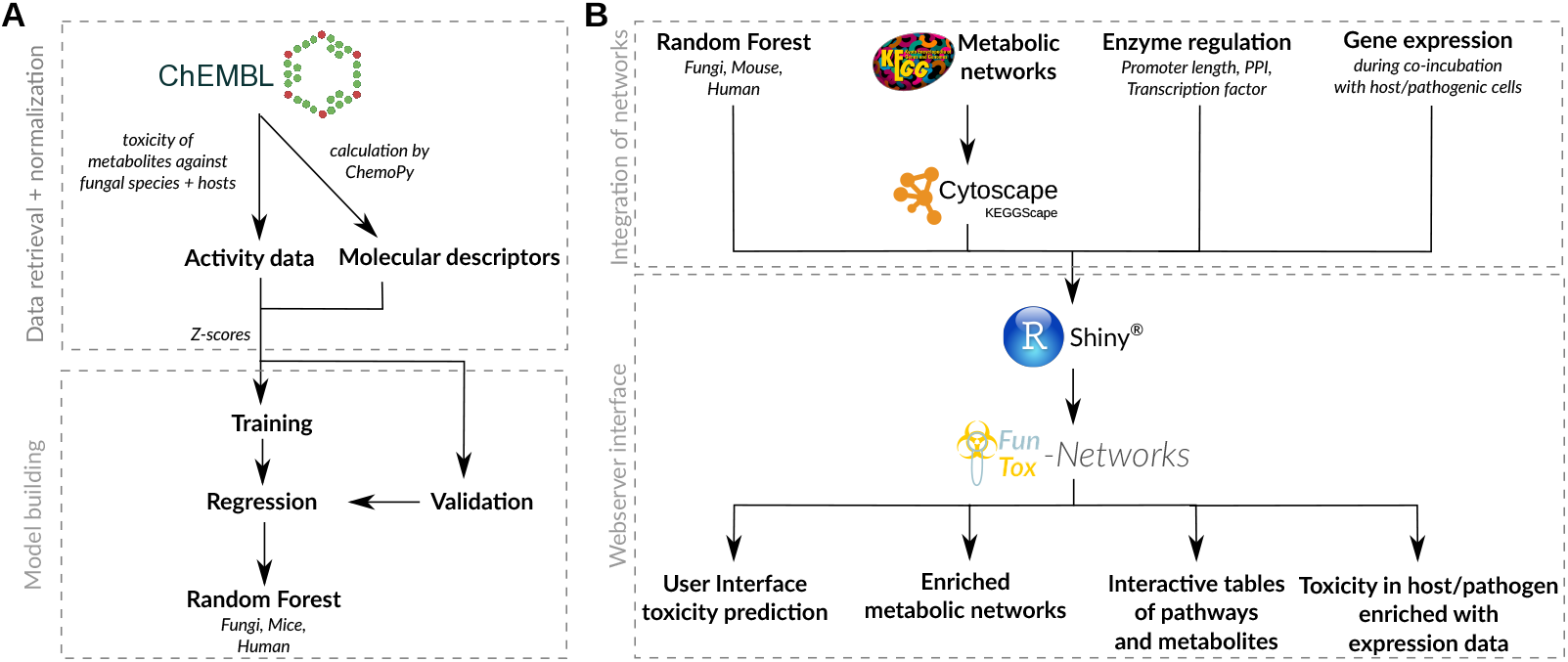
**A** Schematic workflow of data preparation and model building to predict metabolite toxicity in fungi, mice and human. **B** Overview of used databases, data and tools to integrate toxicity prediction and metabolic networks and the provided interfaces in the web-service *FunTox*-*Networks*.

To extract structural features to train a QSAR, we retrieved the Simplified Molecular Input Line Entry Specification (SMILES) [44] for all compounds from the above described data set. Using the python package Mordred [76], we calculated molecular descriptors of several properties such as size, polarity and topology (see Tab. 1). To rule out that descriptor choice weakens prediction performance, multiple sets of descriptors with minimal correlation to each other were tested. However, prediction performance was similar based on this set and we therefore conclude, that our selection of descriptors is sufficient to represent the chemical space.

**TABLE 1.**
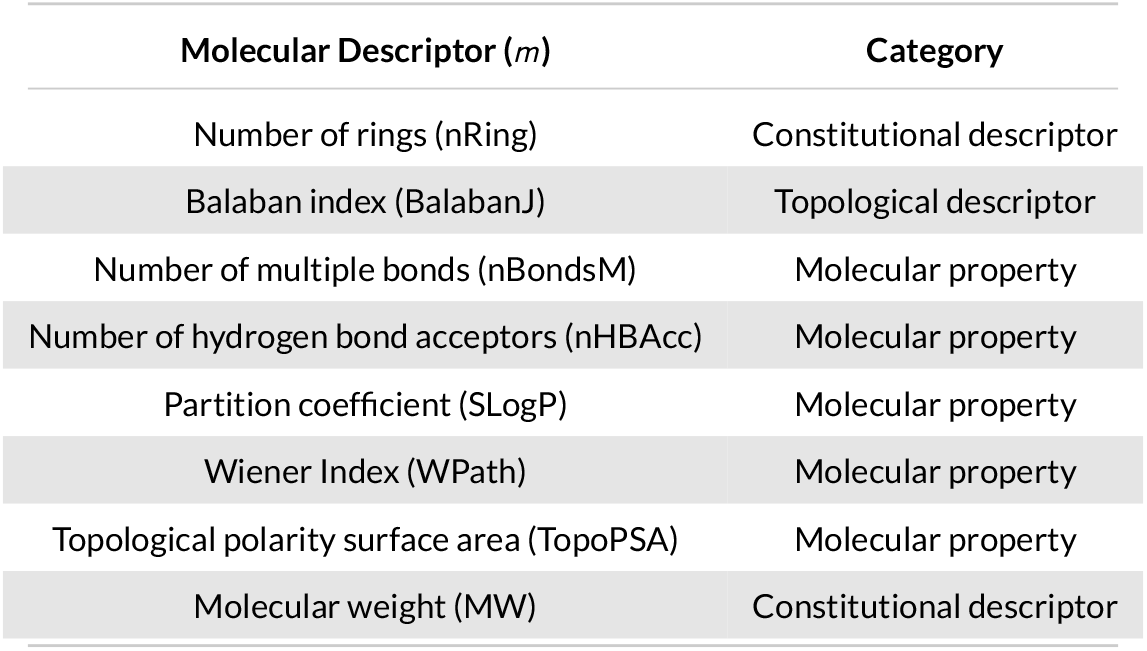
Overview of used molecular descriptors.

In contrast to other QSAR models, we built a multi-output model to train the model on all fungal species as well as on different toxicity measurements (standard types) using a normalization scheme as described previously [37, 38]. Hence, the activity data (*tox*) is standardized by Z-normalization with the mean (*avg*) and standard deviation (*sd*) of all activity data under the same standard type (*s*):

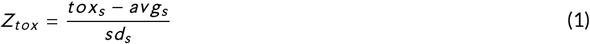

The normalized activity data (*Z_tox_*) is estimated through a regression by structural features (molecular descriptors), which are Z-normalized for fields (*f*) describing the influence of assay characteristics (organism *f_o_*, standard type *f_s_*, cell type *f_c_* and data curation level *f_d_*). Together with the average Z-normalized toxicity for the combination of organism as well as standard type (*Z_exp_*) and the eight molecular descriptors (see Tab. 1) this results in 33 features describing the relationship of toxicity, assay characteristics and molecular structure:

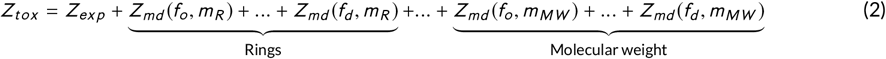

After removal of outliers (e.g. extreme values in ChEMBL for (non-)toxic compounds) based on a simple linear model (upper and lower 5% residuals), a 10 % randomly chosen subset of the resulting data was used to identify the best method for regression by applying the rRegr package [77] comparing several machine learning approaches (see Supp. 6). The decision tree-based random forest regression method [78] performed best in terms of deviation and correlation and therefore random forests were trained on the full data set using the R package ‘ranger’ [79]. As optimal parameters for the machine learning approach, we used five randomly selected features at each split (*mtry*) and one hundred trees (*ntree*), which has proven to be a good trade-off between performance and computational costs in previous machine learning approaches [80] and in for our model (cf. our analysis documented in http://doi.org/10.5281/zenodo.3529162).

Furthermore, cross-validation was performed by splitting the data set in 4 splits with each run using 75% of the full data set as training and 25% as test data set to calculate the cross-validated 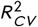 (or *Q* ^2^) [81]. Since values of 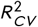 were close to the coefficient of determination *R*^2^, we did not observe overfitting of our model for all splits (cf. our analysis documented in http://doi.org/10.5281/zenodo.3529162) and using all features described in equation 2. Therefore all features were used to train the final random forest models for fungal species, human and mice on the full data set.

For QSAR models it is advised to provide information about the certainty of a prediction, called applicability domain [46]. The robustness of toxicity prediction can be inferred by analyzing the chemical space of compounds used for learning and by the confidence of predictions made by the random forest regression. To this end, the standard deviation across all trees of the random forest, which is an indicator of prediction accuracy [47], is given as additional output. Furthermore, we performed a principle component analysis (PCA) of the molecular descriptor space of all compounds used to train the random forest regression to visualize similarity of queried to trained compounds [82]. Both is provided in the web-service of our toxicity prediction.

### 4.2 Integration of intermediate toxicity and metabolic networks

For the identification of possible drug targets in metabolic networks, pathways of the major fungal pathogens and their hosts were enriched with the predicted toxicity of metabolites and regulatory data of enzymes (see Tab. 2). Pathway maps were retrieved using the R package KEGGREST in the KEGG Markup Language (KGML) format [39, 83] for the organisms listed in Tab. 2. Toxicity of KEGG compounds was predicted with our random forest regression model with MIC (minimal inhibitory concentration) in fungal species as standard type and IC50 (half maximal inhibitory concentration) for human cells respectively LD50 (median lethal dose) for mice, since these were the preferred standard types in the data set for training.

**TABLE 2.** Organisms with toxicity prediction, metabolic networks from KEGG and estimated regulatory effort of enzymes.

| Organism | KEGG ID | Description | Regulatory data |
| --- | --- | --- | --- |
| *Arthroderma benhamiae* | abe | Dermatophyte | Promoter length |
| *Aspergillus fumigatus* | afm | Saprophyte, (invasive) aspergillosis | PPI score |
| *Aspergillus flavus* | afv | Saprophyte and crop pathogen<br>(invasive) aspergillosis | Promoter length |
| *Candida albicans* | cal | Commensal, (invasive) candidiasis | PPI score |
| *Candida glabrata* | cgr | Commensal, (invasive) candidiasis | PPI score |
| *Cryptococcus neoformans* | cnb | Saprophyte, (invasive) cryptococcosis | PPI score |
| *Homo sapiens* | hsa | Host | Transcription factor |
| *Mus musculus* | mmu | Host | Transcription factor |
| *Saccharomyces cerevisiae* | sce | Important fungal model organism<br>rare opportunistic pathogen | PPI score |

We used three different approaches to infer the strength of regulation of individual enzymes referred to as regulatory effort controlling an enzyme. Intuitively, the number of transcription factors controlling an enzyme is a direct measurement of regulatory effort and is used for human and murine enzymes, where genome-wide data is available from the RegNetwork database [84].

For fungi with only scarce knowledge of gene regulatory interactions, we used information that were obtained from the FungiWeb-Database (https://fungiweb.bioapps.biozentrum.uni-wuerzburg.de). Since no experimentally validated large-scale protein-protein networks on a genome-scale for these species exists we used an interolog-based method to infer these networks from the established and validated human and yeast protein-protein networks (see Remmele *et al.* 2015 [85]). For this approach we obtained the intraspecies networks from *Homo sapiens* and *Saccharomyces cerevisiae* from the 14 active partners of the International Molecular Exchange (IMEx) consortium [86] and the protein orthology information from Inparanoid8 [87], *Candida* Genome Database (CGD) [88] and *Aspergillus* Genome Database (AspGD) [89] via an automated pipeline. Based on the experimental evidence, size of the experiment and number of publication that support a source interaction we calculated for each edge of these networks a modified version of the MINT-Score (Molecular INTeraction database [90]) as a measurement of the interaction reliability. From the PPI networks of *A. fumigatus* (176,584 interactions, 4,086 interactors), *C*. *albicans* (97,614 interactions, 1,984 interactors), *C*. *glabrata* (297,419 interactions, 4,604 interactors) and *C. neoformans* (191,274 interactions, 3,203 interactors) we estimated the regulatory effort of an enzyme by the connectivity (vertex degree) in the network of the species and weighted each edge by their evidence according to the reliability score. The resulting PPI score provides a good estimate of regulation and interaction on post-translational level, which we could confirm for yeast (see supplement 8).

For organisms where neither transcription factors nor a PPI score are available (see Tab. 2), the promoter length determined as the intergenic distance was used as in previous studies for prokaryotes [21, 23]. Promoter lengths have proven to be good proxies of regulatory effort due to the reduced genome sizes of fungi compared to higher eukaryotes which makes a loss of non-functional sequences more likely [91]. In consequence, the length of promoter regions should reflect the number of functional elements such as transcription factor binding sites contained in them. In agreement with this hypothesis, we found for the well-studied regulatory network of yeast that promoter length correlates with the number of transcription factors (see Supp. 8).

We enriched pathways maps with color-coded information on regulatory effort and toxicity information in KGML files. We used Cytoscape [92] with the extension KEGGScape [93] to convert KGML formats to JavaScript Object Notation (JSON), which can be used in a javascript-based network viewer provided by Cytoscape. The collection of pathway maps is embedded in the R Shiny [94] application *FunTox*-*Networks* and accessible as a web-service (http://funtox.bioinf.uni-jena.de). Moreover, pathway maps and compounds are summarized in tables that can be searched, filtered and downloaded to facilitate the search of toxic intermediates and the controlling enzymes to identify drug targets.

### 4.3 Investigation of glyoxylate detoxifying enzymes in *C*. *albicans*

#### Identification and co-regulation of enzymes

We used metabolic information of *Saccharomyces cerevisiae* from MetaCyc [95] to identify additional enzymes which degrade glyoxylate. In addition to the malate synthase (EC: 2.3.3.9) this includes the glyoxylate reductase (EC: 1.1.1.79) and the alanine-glyoxylate transaminase (EC: 2.6.1.44). Based on sequence homology to yeast counterparts we determined *GOR1* (orf19.2989), respectively *HBR2* (orf19.1078) and calculated the Spearman’s rank correlation of their gene expression in a published data set of the response in *C*. *albicans’* transcription to organic acids [67].

#### Strains and mutant generation

All *C*. *albicans* strains were stored as glycerol stocks at −80°*C* and streaked on YPD plates for growth at 30°*C* before use. Mutants were generated by standard heat shock transformation procedures (PCR-amplified pFA plasmids with ≈ 100*bp* homology regions [96]) using the BWP17 strain (lacking *URA3*, *HIS1* and *ARG4* [97]). Uridine prototrophy was restored with CIp10, and BWP17 + CIp30 (restored for all three markers) was used as an isogenic control. For additional deletion of *HBR2*, the dominant SAT flipper method was employed as described previously [98], using pSFS5 and the In-Fusion cloning system (Takara Biotech) for creation of a deletion cassette with *Sac*I and *Kpn*I restriction sites. The final mutant was cured of the SAT1 cassette and contained only the FRT site in place of *HBR2*. All primers are listed in supplementary table 3, and all mutants are listed in supplementary table 4.

#### Growth assays and inhibition by glyoxylate

The *C*. *albicans* deletion mutants (*mls1*Δ/Δ, *hbr* 2Δ/Δ, *mls1*Δ/Δ *hbr2*Δ/Δ) or wild type (SC5314) were grown over night in yeast extract peptone dextrose (YPD) medium at 30°*C*, 180 rpm and then washed three times with distilled water. For the assay, 10*μL* of the cell suspension in phosphate-buffered saline (PBS) was mixed (in 96 well plates) with 190*μL* buffered Synthetic Defined (SD) medium (1×YNB, w/o amino acids/ammonium sulfate; 5 *g/L* ammonium sulfate; 2*g/L* glucose; 100*mM* phosphate buffer, *pH* = 3). The medium also contained glyoxylate, in a 1:2 dilution series from 500*mM* down to 7.8*mM*, plus a control without any glyoxylate. The strains were grown in triplicates in each glyoxylate concentration for 50 hours at 30°*C* in a Tecan Infinite 200 microplate reader. The absorption at 600*nm* was measured every 15 minutes after 10*s* of orbital shaking.

#### Long term survival assays

The *C*. *albicans* deletion mutants (*icl1*Δ/Δ, *mls1*Δ/Δ, *hbr* 2Δ/Δ, *mls1*Δ/Δ *hbr2*Δ/Δ) or wild type (SC5314) were grown for long periods in SD medium without carbon source or with 2*g*/*L* of a C2 compound (ethanol). For this, overnight YPD cultures of *C*. *albicans* (30°*C*, 180 rpm) were washed three times with PBS. The cells were counted in a Neubauer chamber, and 20*mL* of each SD medium was inoculated at a concentration of 10^5^ *cells*/*ml*. The strains were grown at 30°*C* for 8 days, and samples (1*mL*) taken every 48 hours for cell number quantification with a Neubauer chamber. A serial 1:10 dilution (down to 10^6^ *cells*/*ml*) in PBS was used to assess the number of living cells in the sample: Three 10*μL* spots of each dilutions were pipetted on YPD agar plates and incubated for 48 hours at 30°*C* for counting of living cells. The plates were then incubated for further 48 hours at 30°*C* to allow growth of smaller colonies from lagging cells, which were added to the count.

#### Measurement of glyoxylate in Candida albicans strains

Overnight pre-cultures of *C*. *albicans* wild type (SC5314) and deletion mutants *mls1*Δ/Δ and *mls1*Δ/Δ *hbr2*Δ/Δ were used to inoculate the corresponding main cultures with an OD600 of 0.1. After a cultivation over a period of 12*h* in 50*mL* YNB broth without amino acids and 2% glucose as carbon source at 30°*C*, samples were harvested in triplicates and intracellular metabolites extracted. For this purpose, the cells were centrifuged at 4°*C* and 4000*g* for 5*min*, followed by three washing steps with 20*mL* cold 0.9% *NaCl* solution. The pelleted cells were resuspended in ethanol supplemented with 2.5*μg /mL* [U-13*C*_5_]-ribitol or [2,2,3,3,4,4-*D*_6_]-glutaric acid as internal standard. Resuspended cells were set out by ultrasonic treatment at 70°*C* for 15*min* for cell lysis. The suspensions were incubated for 2*min* on ice followed by the addition of 0.75*m H*_2_*O*. Metabolites were extracted by addition of 1*mL* chloroform followed by harshly mixing for 1*min* and centrifugation for 5*min* at 4000*g* and 4°*C*. 0.8*mL* of the resulting upper polar phase was sampled and triplicates were dried in a vacuum concentrator over night at 4°*C*. Polar dried metabolites were automatically derivatized in a two step procedure; first, 15*μL* of 2% methoxyamine hydrochloride in pyridine was added and samples were incubated at 55°*C* for 90*min* under shaking. Second, MTBSTFA (N-methyl-N[tert]-butyldimethylsilyl trifluoroacetamide w/1% tert-butyldimethylchlorosilane) was added in an equal amount and samples were incubated for 60*min* at 55°*C*. Gas-chromatography coupled to mass-spectrometry (GC-MS) measurements were performed using an Agilent 7890B GC/Agilent MSD 5977B instrument. Derivatized metabolites were separated by a capillary column (Phenomenex ZB 35, 30*m* length, 0.25*mm* in diameter, 0.25*μm* film thickness) and a SSL Liner Agilent 5190-3171: 900 *μL* (splitless, single taper, wool, Ultra). Carrier gas was Helium with a flow rate of 1.0*mL/mi n*. The temperature in the GC oven was 2*min* at 100°*C* with increased steps of 10°*C* per minute up to 300°*C*. Total run time of 26*min* per sample. Electron impact ionization at 70*eV* were operated by an Agilent MSD 5977B with an extractor source and the transferline temperature at 280°*C*. The mass spectometry was held at 230°*C* and the quadrupol at 150°*C*. The detection of metabolites were done in scan and sim mode. To aquire the full mass spectra scanning from 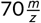 to 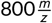 at a scan rate of 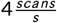. SIM data for glyoxylate was perfomed with the *mz* of 160, 161, 162, 202, 203, 204 each a dwell time of 15*ms*. Data was processed with the MetaboliteDetector software. Statistical analyses were performed. Standard errors of mean (SEM) were shown. Students T-tests were performed and p values are calculated with two way analysis. Significance values were considered as significant with values *p < 0.05, **p < 0.01, ***p < 0.001.

## Acknowledgements

JE and StS acknowledge support by the Deutsche Forschungsgemeinschaft (DFG) within the CRC/Transregio 124 ‘Pathogenic fungi and their human host: Networks of interaction’ (support code 210879364) sub project B1, CL and MD within sub project B2 and BH within sub project C1. BH and PMJ were further supported by the Balance of the Microverse Cluster (Germany’s Excellence Strategy – EXC 2051 – Project-ID 390713860). CK acknowledges support by the Cluster of Excellence ‘Precision Medicine in Chronic Inflammation’ (support code EXC 2167). HGD thanks support of Ministry of Science and Innovation MICIIN (PID2019-104148GB-I00) and Basque Government (IT1045-16). We thank Elina Wiechens for programming parts of the prediction pipeline.

## Supporting Information

Data and scripts of machine learning pipeline are documented and stored here: http://doi.org/10.5281/zenodo.3529162.

### Details of machine learning procedure

#### Performance of regression models

**FIGURE 6.**
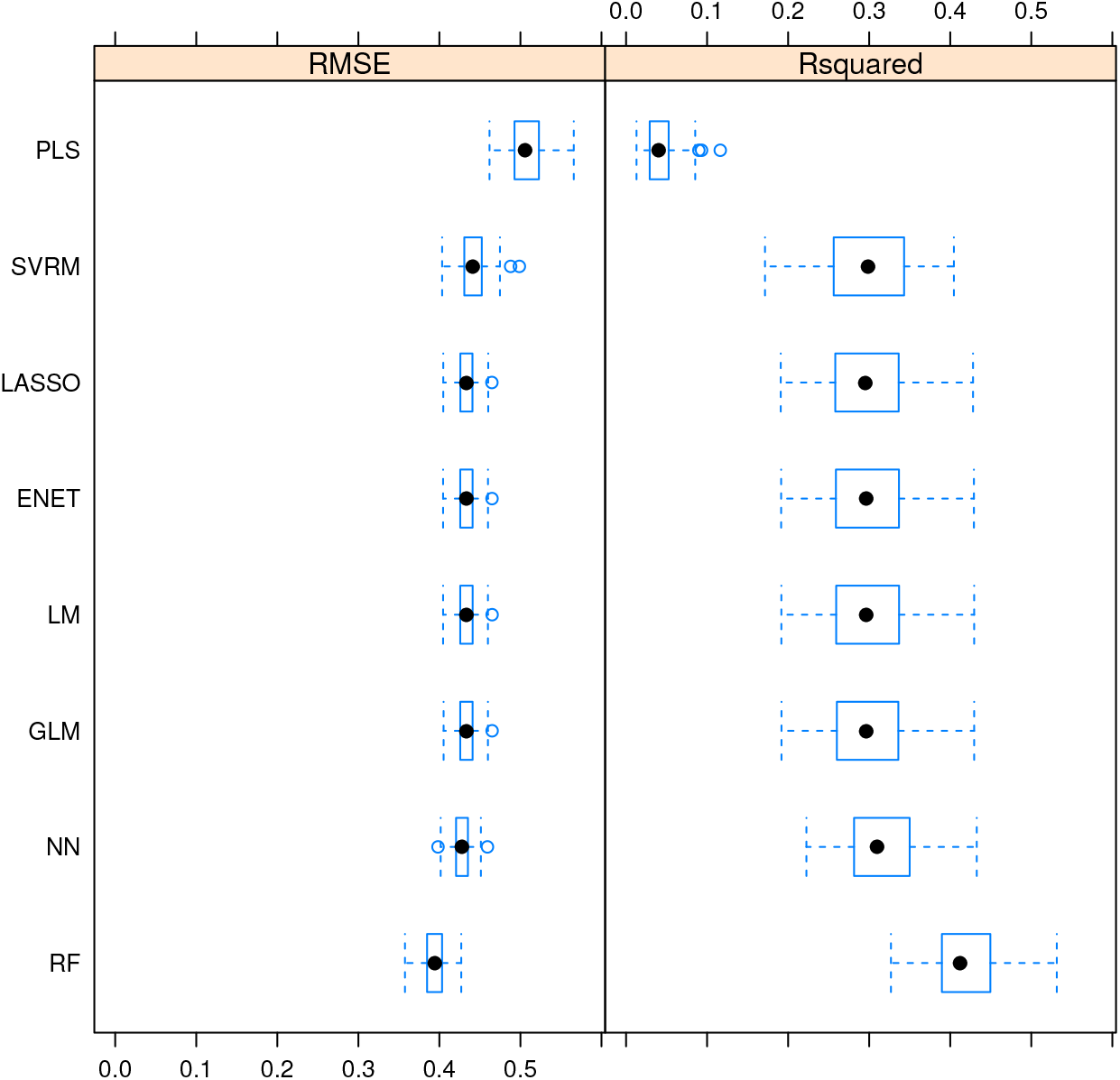
Training results of different regression models measured by the root mean squared error (RMSE) and the coefficient of determination (*R*^2^). Models: Multiple Linear regression (LM), Generalized Linear Model with Stepwise Feature Selection (GLM), Partial Least Squares Regression (PLS), Lasso regression (LASSO), Elastic Net regression (ENET), Support vector machine using radial functions (SVRM), Neural Networks regression (NN), Random Forest (RF).

**FIGURE 7.**
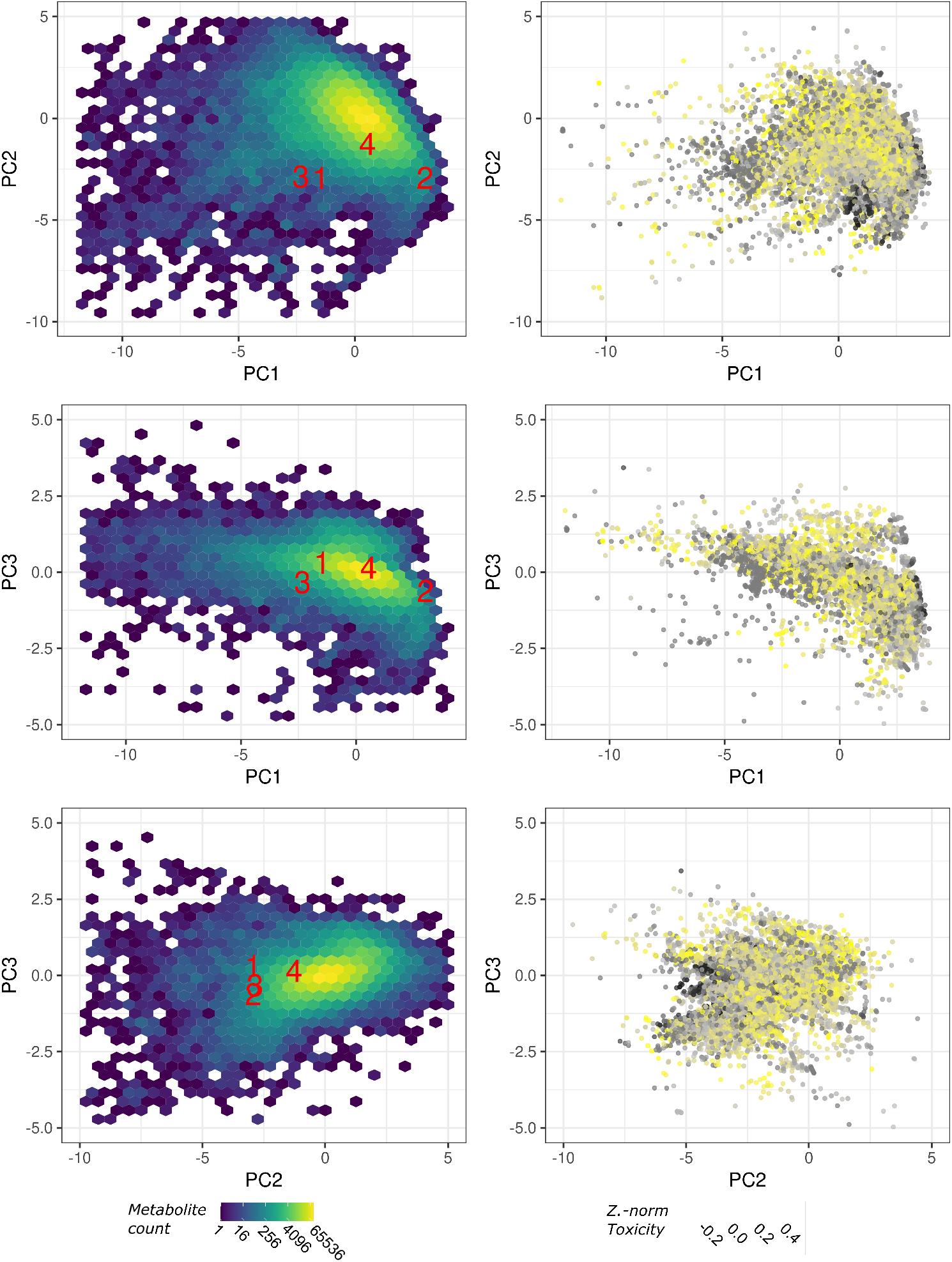
Additional plots including third principle component for applicability domain (left) and toxicity distribution (right) of metabolites as shown in

### Regulatory effort estimates in yeast

**FIGURE 8.**
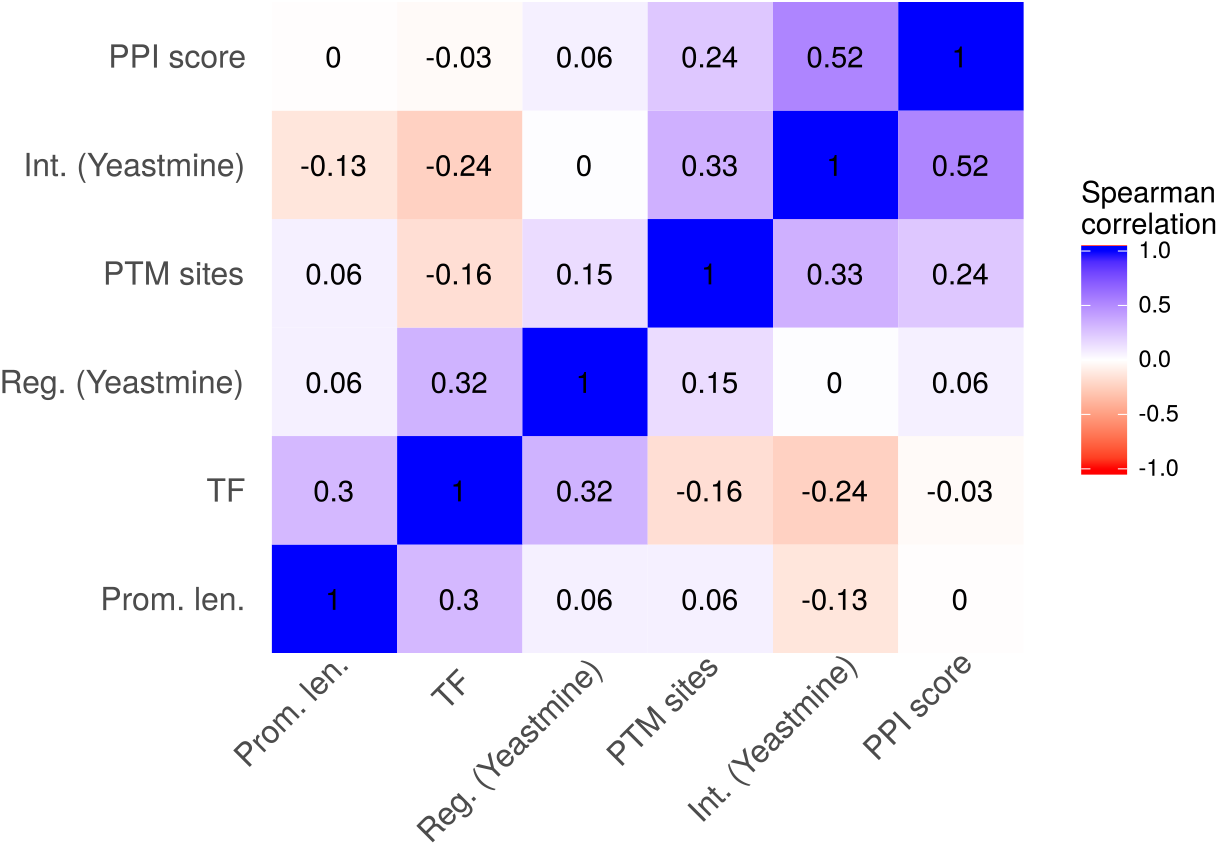
Correlation of promoter length, number of transcription factors (TF) from Yeastract [99], number of regulators as well as interactions listed in YeastMine [100], number of PTM sites from dbPTM [101] and the PPI score for each gene in yeast.

### Primer and mutant list

**TABLE 3.**
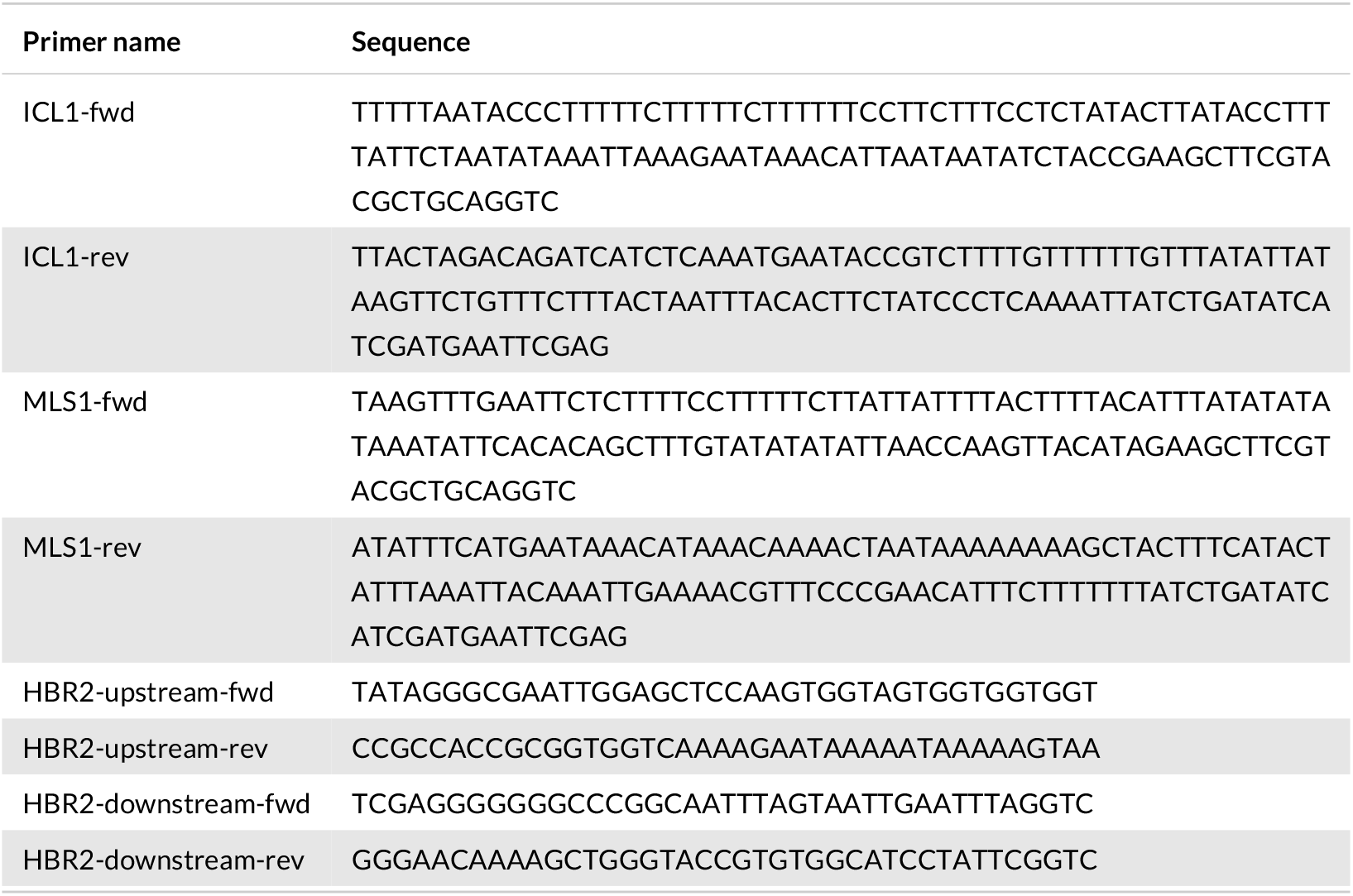
List of primers

**TABLE 4.**
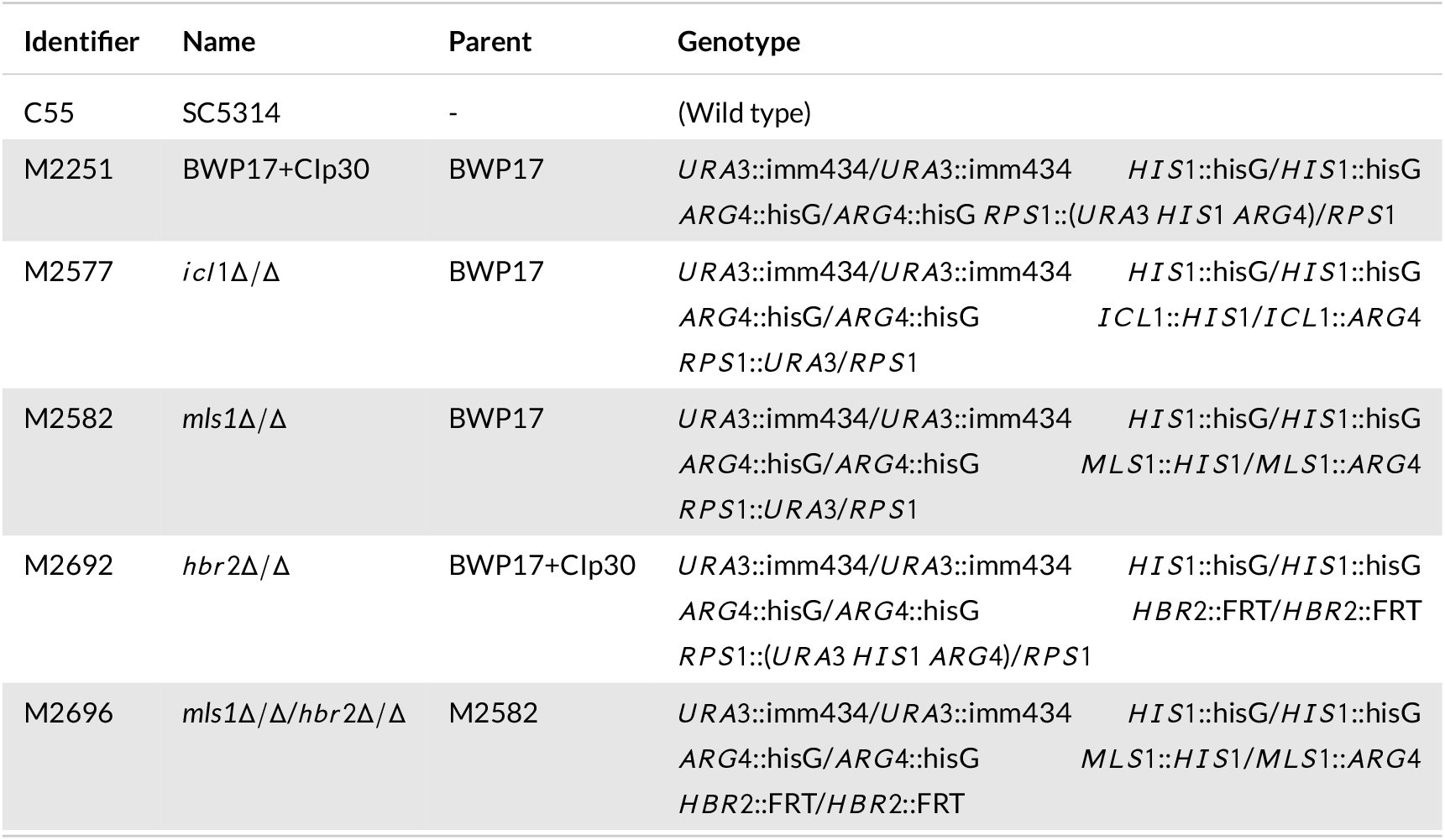
List of strains and mutants

## Notes

http://funtox.bioinf.uni-jena.de

http://doi.org/10.5281/zenodo.3529162

### Competing Interest Statement

The authors have declared no competing interest.

## REFERENCES

[1] Havlickova B, Czaika VA, Friedrich M. Epidemiological trends in skin mycoses worldwide. Mycoses 2008;51(s4):2–15.

[2] Brown GD, Denning DW, Gow NA, Levitz SM, Netea MG, White TC. Hidden killers: human fungal infections. Science Translational Medicine 2012;4(165):165rv13–165rv13.

[3] Gangneux JP, Bougnoux ME, Hennequin C, Godet C, Chandenier J, Denning D, et al. An estimation of burden of serious fungal infections in France. Journal de Mycologie Medicale 2016;26(4):385–390.

[4] Hoenigl M. Invasive Fungal Disease Complicating Coronavirus Disease 2019: When It Rains, It Spores. Clinical Infectious Diseases 2020;.

[5] Raut A, Huy NT. Rising incidence of mucormycosis in patients with COVID-19: another challenge for India amidst the second wave? The Lancet Respiratory Medicine 2021;.

[6] Low CY, Rotstein C. Emerging fungal infections in immunocompromised patients. F1000 Medicine Reports 2011;3.

[7] Enoch DA, Yang H, Aliyu SH, Micallef C. The changing epidemiology of invasive fungal infections. Human Fungal Pathogen Identification 2017;p. 17–65.

[8] Roemer T, Krysan DJ. Antifungal drug development: challenges, unmet clinical needs, and new approaches. Cold Spring Harbor perspectives in medicine 2014;4(5):a019703.

[9] Ford CB, Funt JM, Abbey D, Issi L, Guiducci C, Martinez DA, et al. The evolution of drug resistance in clinical isolates of *Candida albicans*. eLife 2015;4:e00662.

[10] Van Daele R, Spriet I, Wauters J, Maertens J, Mercier T, Van Hecke S, et al. Antifungal drugs: what brings the future? Medical Mycology 2019;57(Supplement_3):S328–S343.

[11] Wall G, Lopez-Ribot JL. Current antimycotics, new prospects, and future approaches to antifungal therapy. Antibiotics 2020;9(8):445.

[12] Ene IV, Brunke S, Brown AJ, Hube B. Metabolism in fungal pathogenesis. Cold Spring Harbor Perspectives in Medicine 2014;4(12):a019695.

[13] K Mazu T, A Bricker B, Flores-Rozas H, Y Ablordeppey S. The Mechanistic Targets of Antifungal Agents: An Overview. Mini Reviews in Medicinal Chemistry 2016;16(7):555–578.

[14] McCarthy MW, Walsh TJ. Amino acid metabolism and transport mechanisms as potential antifungal targets. International Journal of Molecular Sciences 2018;19(3):909.

[15] Sun S, Chen X, Zhang Z, Chen Z, Li Y, Su S. Potential antifungal targets based on glucose metabolism pathways of *Candida albicans*. Frontiers in Microbiology 2020;11:296.

[16] Kämmer P, McNamara S, Wolf T, Conrad T, Allert S, Gerwien F, et al. Survival Strategies of Pathogenic Candida Species in Human Blood Show Independent and Specific Adaptations. mBio 2020;11(5).

[17] Ewald J, Sieber P, Garde R, Lang SN, Schuster S, Ibrahim B. Trends in mathematical modeling of host–pathogen interactions. Cellular and Molecular Life Sciences 2020;77(3):467–480.

[18] Klipp E, Heinrich R, Holzhütter HG. Prediction of temporal gene expression. European Journal of Biochemistry 2002;269(22):5406–5413.

[19] Wessely F, Bartl M, Guthke R, Li P, Schuster S, Kaleta C. Optimal regulatory strategies for metabolic pathways in *Escherichia coli* depending on protein costs. Molecular Systems Biology 2011;7(1):515.

[20] Bartl M, Kötzing M, Schuster S, Li P, Kaleta C. Dynamic optimization identifies optimal programmes for pathway regulation in prokaryotes. Nature Communications 2013;4.

[21] de Hijas-Liste GM, Balsa-Canto E, Ewald J, Bartl M, Li P, Banga JR, et al. Optimal programs of pathway control: dissecting the influence of pathway topology and feedback inhibition on pathway regulation. BMC Bioinformatics 2015;16(1):163.

[22] Ewald J, Kötzing M, Bartl M, Kaleta C. Footprints of optimal protein assembly strategies in the operonic structure of prokaryotes. Metabolites 2015;5(2):252–269.

[23] Ewald J, Bartl M, Dandekar T, Kaleta C. Optimality principles reveal a complex interplay of intermediate toxicity and kinetic efficiency in the regulation of prokaryotic metabolism. PLoS Computational Biology 2017;13(2):e1005371.

[24] Ewald J, Bartl M, Kaleta C. Deciphering the regulation of metabolism with dynamic optimization: an overview of recent advances. Biochemical Society Transactions 2017;.

[25] Kaltdorf M, Srivastava M, Gupta SK, Liang C, Binder J, Dietl AM, et al. Systematic identification of anti-fungal drug targets by a metabolic network approach. Frontiers in Molecular Biosciences 2016;3.

[26] Kemble H, Eisenhauer C, Couce A, Chapron A, Magnan M, Gautier G, et al. Flux, toxicity, and expression costs generate complex genetic interactions in a metabolic pathway. Science Advances 2020;6(23):eabb2236.

[27] Liu D, Mannan AA, Han Y, Oyarzún DA, Zhang F. Dynamic metabolic control: towards precision engineering of metabolism. Journal of Industrial Microbiology and Biotechnology 2018;45(7):535–543.

[28] Lee N, Spears ME, Carlisle AE, Kim D. Endogenous toxic metabolites and implications in cancer therapy. Oncogene 2020;39(35):5709–5720.

[29] Santana L, Uriarte E, González-Díaz H, Zagotto G, Soto-Otero R, Méndez-Álvarez E. A QSAR model for in silico screening of MAO-A inhibitors. Prediction, synthesis, and biological assay of novel coumarins. Journal of Medicinal Chemistry 2006;49(3):1149–1156.

[30] Verma J, Khedkar VM, Coutinho EC. 3D-QSAR in drug design-a review. Current Topics in Medicinal Chemistry 2010;10(1):95–115.

[31] Carbonell P, Planson AG, Fichera D, Faulon JL. A retrosynthetic biology approach to metabolic pathway design for therapeutic production. BMC Systems Biology 2011;5(1):122.

[32] Planson AG, Carbonell P, Paillard E, Pollet N, Faulon JL. Compound toxicity screening and structure–activity relationship modeling in *Escherichia coli*. Biotechnology and Bioengineering 2012;109(3):846–850.

[33] Raies AB, Bajic VB. In silico toxicology: computational methods for the prediction of chemical toxicity. Wiley Interdisciplinary Reviews: Computational Molecular Science 2016;6(2):147–172.

[34] Roy K, Kar S, Das RN. Understanding the basics of QSAR for applications in pharmaceutical sciences and risk assessment. Academic press; 2015.

[35] Bento AP, Gaulton A, Hersey A, Bellis LJ, Chambers J, Davies M, et al. The ChEMBL bioactivity database: an update. Nucleic Acids Research 2014;42(D1):D1083–D1090.

[36] Prado-Prado FJ, Borges F, Perez-Montoto LG, González-Díaz H. Multi-target spectral moment: QSAR for antifungal drugs vs. different fungi species. European Journal of Medicinal Chemistry 2009;44(10):4051–4056.

[37] M Casañola-Martin G, Le-Thi-Thu H, Pérez-Giménez F, Marrero-Ponce Y, Merino-Sanjuán M, Abad C, et al. Multi-output Model with Box-Jenkins Operators of Quadratic Indices for Prediction of Malaria and Cancer Inhibitors Targeting Ubiquitin-Proteasome Pathway (UPP) Proteins. Current Protein and Peptide Science 2016;17(3):220–227.

[38] Romero-Durán FJ, Alonso N, Yañez M, Caamaño O, García-Mera X, González-Díaz H. Brain-inspired cheminformatics of drug-target brain interactome, synthesis, and assay of TVP1022 derivatives. Neuropharmacology 2016;103:270–278.

[39] Kanehisa M, Furumichi M, Tanabe M, Sato Y, Morishima K. KEGG: new perspectives on genomes, pathways, diseases and drugs. Nucleic Acids Research 2017;45(D1):D353–D361.

[40] Alexander D, Tropsha A, Winkler DA. Beware of R 2: simple, unambiguous assessment of the prediction accuracy of QSAR and QSPR models. Journal of Chemical Information and Modeling 2015;55(7):1316–1322.

[41] Wang C, Wei Z, Wang L, Sun P, Wang Z. Assessment of bromide-based ionic liquid toxicity toward aquatic organisms and QSAR analysis. Ecotoxicology and Environmental Safety 2015;115:112–118.

[42] Roberts MC. Antibiotic toxicity, interactions and resistance development. Periodontology 2000 2002;28(1):280–297.

[43] Rolain J, Baquero F. The refusal of the Society to accept antibiotic toxicity: missing opportunities for therapy of severe infections. Clinical Microbiology and Infection 2016;22(5):423–427.

[44] Weininger D, Weininger A, Weininger JL. SMILES. 2. Algorithm for generation of unique SMILES notation. Journal of Chemical Information and Computer Sciences 1989;29(2):97–101.

[45] Heller S, McNaught A, Stein S, Tchekhovskoi D, Pletnev I. InChI-the worldwide chemical structure identifier standard. Journal of Cheminformatics 2013;5(1):7.

[46] OECD. Guidance Document on the Validation of (Quantitative) Structure-Activity Relationship [(Q)SAR] Models. Organisation for Economic Co-operation and Development: Paris, France 2007;http://dx.doi.org/10.1787/9789264085442-en.

[47] Sheridan RP. Three useful dimensions for domain applicability in QSAR models using random forest. Journal of Chemical Information and Modeling 2012;52(3):814–823.

[48] Allaman I, Bélanger M, Magistretti PJ. Methylglyoxal, the dark side of glycolysis. Frontiers in Neuroscience 2015;9:23.

[49] Kumar S, Bandyopadhyay U. Free heme toxicity and its detoxification systems in human. Toxicology Letters 2005;157(3):175–188.

[50] Brass EP. Overview of coenzyme A metabolism and its role in cellular toxicity. Chemico-Biological Interactions 1994;90(3):203–214.

[51] Lastauskienė E, Zinkevičienė A, Girkontaitė I, Kaunietis A, Kvedarieneė V. Formic acid and acetic acid induce a programmed cell death in pathogenic *Candida* species. Current Microbiology 2014;69(3):303–310.

[52] Tucey TM, Verma J, Harrison PF, Snelgrove SL, Lo TL, Scherer AK, et al. Glucose homeostasis is important for immune cell viability during Candida challenge and host survival of systemic fungal infection. Cell Metabolism 2018;27(5):988–1006.

[53] Watkins TN, Liu H, Chung M, Hazen TH, Hotopp JCD, Filler SG, et al. Comparative transcriptomics of *Aspergillus fumigatus* strains upon exposure to human airway epithelial cells. Microbial Genomics 2018;4(2).

[54] Kontoyiannis DP. Modulation of fluconazole sensitivity by the interaction of mitochondria and erg3p in *Saccharomyces cerevisiae*. Journal of Antimicrobial Chemotherapy 2000;46(2):191–197.

[55] Richardson MD. Opportunistic and pathogenic fungi. Journal of Antimicrobial Chemotherapy 1991;28(suppl_A):1–11.

[56] Xu S, Shinohara ML. Tissue-resident macrophages in fungal infections. Frontiers in Immunology 2017;8:1798.

[57] Jastrzebowska K, Gabriel I. Inhibitors of amino acids biosynthesis as antifungal agents. Amino Acids 2015;47(2):227–249.

[58] Suliman HS, Appling DR, Robertus JD. The gene for cobalamin-independent methionine synthase is essential in *Candida albicans*: a potential antifungal target. Archives of Biochemistry and Biophysics 2007;467(2):218–226.

[59] Amich J, Krappmann S. Deciphering metabolic traits of the fungal pathogen *Aspergillus fumigatus*: redundancy vs. essentiality. Frontiers in Microbiology 2012;3:414.

[60] Ries LNA, Beattie S, Cramer RA, Goldman GH. Overview of carbon and nitrogen catabolite metabolism in the virulence of human pathogenic fungi. Molecular Microbiology 2018;107(3):277–297.

[61] Amich J, Dümig M, O’Keeffe G, Binder J, Doyle S, Beilhack A, et al. Exploration of sulfur assimilation of *Aspergillus fumigatus* reveals biosynthesis of sulfur-containing amino acids as a virulence determinant. Infection and Immunity 2016;84(4):917–929.

[62] Chebaro Y, Lorenz M, Fa A, Zheng R, Gustin M. Adaptation of *Candida albicans* to Reactive Sulfur Species. Genetics 2017;206(1):151–162.

[63] del Río LA, Corpas FJ, Sandalio LM, Palma JM, Gómez M, Barroso JB. Reactive oxygen species, antioxidant systems and nitric oxide in peroxisomes. Journal of Experimental Botany 2002;53(372):1255–1272.

[64] Lorenz MC, Fink GR. The glyoxylate cycle is required for fungal virulence. Nature 2001;412(6842):83.

[65] Kitahara N, Morisaka H, Aoki W, Takeda Y, Shibasaki S, Kuroda K, et al. Description of the interaction between *Candida albicans*. AMB Express 2015;5(1):1–12.

[66] Puckett S, Trujillo C, Wang Z, Eoh H, Ioerger TR, Krieger I, et al. Glyoxylate detoxification is an essential function of malate synthase required for carbon assimilation in *Mycobacterium tuberculosis*. Proceedings of the National Academy of Sciences 2017;114(11):E2225–E2232.

[67] Cottier F, Tan ASM, Chen J, Lum J, Zolezzi F, Poidinger M, et al. The transcriptional stress response of *Candida albicans* to weak organic acids. G3: Genes, Genomes, Genetics 2015;5(4):497–505.

[68] Schlösser T, Gätgens C, Weber U, Stahmann KP. Alanine: glyoxylate aminotransferase of *Saccharomyces cerevisiae*– encoding gene AGX1 and metabolic significance. Yeast 2004;21(1):63–73.

[69] Miramón P, Lorenz MC. A feast for Candida: metabolic plasticity confers an edge for virulence. PLoS Pathogens 2017;13(2):e1006144.

[70] Cheah HL, Lim V, Sandai D. Inhibitors of the glyoxylate cycle enzyme ICL1 in *Candida albicans* for potential use as anti-fungal agents. PloS One 2014;9(4):e95951.

[71] Fisher MC, Henk DA, Briggs CJ, Brownstein JS, Madoff LC, McCraw SL, et al. Emerging fungal threats to animal, plant and ecosystem health. Nature 2012;484(7393):186.

[72] Dahl RH, Zhang F, Alonso-Gutierrez J, Baidoo E, Batth TS, Redding-Johanson AM, et al. Engineering dynamic pathway regulation using stress-response promoters. Nature Biotechnology 2013;31(11):1039.

[73] Keasling JD. Manufacturing molecules through metabolic engineering. Science 2010;330(6009):1355–1358.

[74] Koivistoinen OM, Kuivanen J, Barth D, Turkia H, Pitkänen JP, Penttilä M, et al. Glycolic acid production in the engineered yeasts *Saccharomyces cerevisiae* and *Kluyveromyces lactis*. Microbial Cell Factories 2013;12(1):82.

[75] Salusjärvi L, Havukainen S, Koivistoinen O, Toivari M. Biotechnological production of glycolic acid and ethylene glycol: current state and perspectives. Applied Microbiology and Biotechnology 2019;103(6):2525–2535.

[76] Moriwaki H, Tian YS, Kawashita N, Takagi T. Mordred: a molecular descriptor calculator. Journal of Cheminformatics 2018;10(1):1–14.

[77] Tsiliki G, Munteanu CR, Seoane JA, Fernandez-Lozano C, Sarimveis H, Willighagen EL. RRegrs: an R package for computer-aided model selection with multiple regression models. Journal of Cheminformatics 2015;7(1):46.

[78] Breiman L. Random forests. Machine Learning 2001;45(1):5–32.

[79] Wright MN, Ziegler A. ranger: A fast implementation of random forests for high dimensional data in C++ and R. arXiv preprint arXiv:150804409 2015;.

[80] Oshiro TM, Perez PS, Baranauskas JA. How Many Trees in a Random Forest? In: International Workshop on Machine Learning and Data Mining in Pattern Recognition Springer; 2012. p. 154–168.

[81] Consonni V, Ballabio D, Todeschini R. Comments on the definition of the Q2 parameter for QSAR validation. Journal of Chemical Information and Modeling 2009;49(7):1669–1678.

[82] Sahigara F, Mansouri K, Ballabio D, Mauri A, Consonni V, Todeschini R. Comparison of different approaches to define the applicability domain of QSAR models. Molecules 2012;17(5):4791–4810.

[83] Tenenbaum D. KEGGREST: Client-side REST access to KEGG; 2017, r package version 1.16.0.

[84] Liu ZP, Wu C, Miao H, Wu H. RegNetwork: an integrated database of transcriptional and post-transcriptional regulatory networks in human and mouse. Database 2015;2015:bav095.

[85] Remmele CW, Luther CH, Balkenhol J, Dandekar T, Müller T, Dittrich MT. Integrated inference and evaluation of host– fungi interaction networks. Frontiers in Microbiology 2015;6:764.

[86] Orchard S, Kerrien S, Abbani S, Aranda B, Bhate J, Bidwell S, et al. Protein interaction data curation: the International Molecular Exchange (IMEx) consortium. Nature Methods 2012;9(4):345.

[87] Sonnhammer EL, Östlund G. InParanoid 8: orthology analysis between 273 proteomes, mostly eukaryotic. Nucleic Acids Research 2014;43(D1):D234–D239.

[88] Arnaud MB, Costanzo MC, Skrzypek MS, Binkley G, Lane C, Miyasato SR, et al. The Candida Genome Database (CGD), a community resource for *Candida albicans* gene and protein information. Nucleic Acids Research 2005;33(suppl_1):D358–D363.

[89] Cerqueira GC, Arnaud MB, Inglis DO, Skrzypek MS, Binkley G, Simison M, et al. The Aspergillus Genome Database: multispecies curation and incorporation of RNA-Seq data to improve structural gene annotations. Nucleic Acids Research 2013;42(D1):D705–D710.

[90] Licata L, Briganti L, Peluso D, Perfetto L, Iannuccelli M, Galeota E, et al. MINT, the molecular interaction database: 2012 update. Nucleic Acids Research 2012;40(D1):D857–D861.

[91] Noble LM, Andrianopoulos A. Fungal genes in context: genome architecture reflects regulatory complexity and function. Genome Biology and Evolution 2013;5(7):1336–1352.

[92] Shannon P, Markiel A, Ozier O, Baliga NS, Wang JT, Ramage D, et al. Cytoscape: a software environment for integrated models of biomolecular interaction networks. Genome Research 2003;13(11):2498–2504.

[93] Nishida K, Ono K, Kanaya S, Takahashi K. KEGGscape: a Cytoscape app for pathway data integration. F1000 Research 2014;3.

[94] Chang W, Cheng J, Allaire J, Xie Y, McPherson J. shiny: Web Application Framework for R; 2017, https://CRAN.R-project.org/package=shiny, r package version 1.0.3.

[95] Caspi R, Billington R, Fulcher CA, Keseler IM, Kothari A, Krummenacker M, et al. The MetaCyc database of metabolic pathways and enzymes. Nucleic Acids Research 2017;46(D1):D633–D639.

[96] Gola S, Martin R, Walther A, Dünkler A, Wendland J. New modules for PCR-based gene targeting in *Candida albicans*: rapid and efficient gene targeting using 100 bp of flanking homology region. Yeast 2003;20(16):1339–1347.

[97] Wilson RB, Davis D, Mitchell AP. Rapid hypothesis testing with *Candida albicans* through gene disruption with short homology regions. Journal of Bacteriology 1999;181(6):1868–1874.

[98] Sasse C, Schillig R, Dierolf F, Weyler M, Schneider S, Mogavero S, et al. The transcription factor Ndt80 does not contribute to Mrr1-, Tac1-, and Upc2-mediated fluconazole resistance in *Candida albicans*. PloS One 2011;6(9):e25623.

[99] Teixeira MC, Monteiro PT, Guerreiro JF, Gonçalves JP, Mira NP, dos Santos SC, et al. The YEASTRACT database: an upgraded information system for the analysis of gene and genomic transcription regulation in *Saccharomyces cerevisiae*. Nucleic Acids Research 2013;42(D1):D161–D166.

[100] Balakrishnan R, Park J, Karra K, Hitz BC, Binkley G, Hong EL, et al. YeastMine—an integrated data warehouse for *Saccharomyces cerevisiae* data as a multipurpose tool-kit. Database 2012;2012.

[101] Huang KY, Su MG, Kao HJ, Hsieh YC, Jhong JH, Cheng KH, et al. dbPTM 2016: 10-year anniversary of a resource for post-translational modification of proteins. Nucleic Acids Research 2015;44(D1):D435–D446.

